# Multiomics Approach to Understanding Olaparib Resistance and Predicting Drug Response

**DOI:** 10.1101/2023.04.04.535542

**Authors:** Won-Jun Lim, Hyunjin M. Kim, YongHo Oh, Junhee Pyo

## Abstract

We aimed to uncover genetic factors affecting resistance to the cancer drug olaparib. To do this, we utilized multiomics matrix factorization (MOFA), a multiomics approach, to explore omic-based features that might become biomarker candidates. Our results showed that 17 damaging mutations, 6 gene expression signatures, 17 DNA methylations, and 26 transcription-factor activities can impact the refractory response to olaparib.

To verify the potential utility of the identified biomarker candidates, we generated a predictive model to differentiate between olaparib responding and nonresponding cell lines using machine learning techniques, including support vector machine algorithms, random forest algorithms, and Siamese neural networks. The model was centered around six gene-expression biomarker candidates and validated using the Genomics of Drug Sensitivity in Cancer database.

Our findings suggest that using a multiomics approach with machine learning methods can lead to a better understanding of the mechanism of drug resistance and identify biomarkers, which will ultimately facilitate the appropriate administration of drugs to patients. The source codes can be found at https://github.com/wjlim/DrugResistance.

## Introduction

Olaparib is a Poly ADP-Ribose Polymerase (PARP) inhibitor that has shown promising results in the treatment of various types of cancers. It has been particularly effective in the treatment of malignancies in patients with BRCA1/2 mutations (1). PARP is an enzyme involved in DNA repair, and its inhibition by olaparib leads to the accumulation of DNA damage and subsequent cell death in cancer cells (2). Several clinical trials have investigated the safety and efficacy of olaparib in different types of cancer, including ovarian, breast, prostate, and pancreatic cancers. In 2014, olaparib became the first PARP inhibitor to receive approval from the US Food and Drug Administration (FDA) for the treatment of advanced ovarian cancer in patients with BRCA1/2 mutations. Since then, olaparib has been approved for the treatment of several other types of cancers. Despite its potential therapeutic benefits, olaparib has also been associated with adverse effects, including anemia, fatigue, and gastrointestinal symptoms (3). Moreover, resistance to olaparib has been observed in some patients, highlighting the need for a better understanding of the mechanisms underlying resistance and the development of strategies to overcome it (4). Given the promising results and potential limitations of olaparib, further research is needed to explore its efficacy in different types of cancer, identify predictive biomarkers of response, and develop novel strategies to overcome resistance.

Biomarkers can be used to detect and monitor disease progression, therapeutic response, and drug resistance (5,6). They can provide crucial information about the underlying mechanisms of drug resistance and help to develop new strategies for effective treatment (7). In the context of drug resistance research, biomarkers can help to identify patients who are at risk of developing resistance to specific drugs, predict the likelihood of resistance, and monitor the progression of resistance over time (7,8). This information can be used to personalize treatment, tailor therapy to the individual patient, and increase the effectiveness of treatments. By understanding the molecular mechanisms of drug resistance and identifying specific biomarkers, researchers can develop new drugs and treatment regimens that are better suited to target resistant cells, reduce the development of resistance, and improve patient outcomes.

In the discovery of biomarkers, there are numerous limitations that need to be addressed. One major challenge is that current biomarkers often have inadequate specificity and sensitivity (9,10). Many biomarkers are not able to accurately distinguish between disease and nondisease states, leading to false positive or false negative results. Another limitation is the lack of reproducibility in the results obtained from different studies using the same biomarker. This inconsistency can be attributed to a variety of factors, including differences in patient populations, sample preparation, and analytical methods.

Nevertheless, the identification of biomarkers can provide valuable insights into various diseases and help to develop effective diagnostic and therapeutic strategies. Machine learning and deep learning are two powerful approaches that have been increasingly applied to the discovery of biomarkers (11).

Few-shot learning is a deep learning approach that is particularly useful in drug resistance research (12,13). This approach is designed to address the problem of limited data availability, which is a common challenge in biomarker discovery. In traditional deep learning, large amounts of labeled data are required to train the model effectively. However, the availability of labeled, annotated data relevant to drug resistance is often limited. Few-shot learning offers a solution to limited data (14). The method uses prior knowledge from related tasks, as well as transfer learning techniques, which enable the model to learn new tasks with limited data. This approach can allow models to generalize well with limited training data (15,16).

In this study we investigated cancer cell lines derived from 38 patients in the cancer cell line encyclopedia (CCLE) database to identify genetic features associated with resistance to olaparib and biomarker candidates. Additionally, the performance of these features was tested using machine learning methods.

## Materials and Methods

### Data Resources

We selected 19 responders to olaparib and 33 nonresponders from the CCLE based on the criteria of olaparib responders and nonresponders defined in 206162Orig1s000 (17). The cell lines were chosen based on the availability of RNA expression data, which were obtained from the DepMap portal (DepMap, Broad Institute (2022) DepMap 22Q1 Public figshare dataset: https://doi.org/10.6084/m9.figshare.19139906.v1). Our analysis included data on damaging mutations, RNA expression, and DNA methylation, which were obtained from RRBS data among the larger set of data available in the CCLE database, including damaging mutations, RNA expression, fusion calls, DNA methylation by RRBS, histone H3 modification, proteomics, and metabolomics. We used 19,177 transcripts, 17,313 hotspot mutations, and 51,845 CGI methylations from DepMap for our analysis.

### Multiomics Data Analysis

We utilized multiomics factor analysis v2 (MOFA+, v2_1.2.2) to identify the most significant features that differentiate responders from nonresponders (18). MOFA provides a more comprehensive functional inference through gene-set enrichment analysis than gene expression count data due to its use of matrix factorization to find latent vectors of the each features and its explanation of data modalities through variance decomposition (19). To apply this knowledge to our dataset, we employed DoRothEA to infer protein and pathway activities (20). DoRothEA annotates and measures protein activities by utilizing gene networks that interact with transcription factors. We selected genes with high-confidence evidence to predict protein activity and generate the matrix, using curated databases, ChIP-seq, transcription factor binding sites (TFBS), and inferred genotype-tissue expression (GTEx) as the sources of evidence. In total, 289 transcription factors with a medium or higher level of confidence, with two or more types of evidence, were used.

### Data Preprocessing

We filtered the significantly changed features to remove the highly variable features and retain only the most important features, as per the guidelines on the MOFA2 website (https://biofam.github.io/MOFA2/faq.html). The edgeR software (version 3.34.1) was utilized to calculate differentially expressed genes (21). Among the 19,177 transcripts, 4,327 genes with significant changes (p <= 0.05) were selected for analysis. A Wilcoxon rank-sum test was performed to identify the most significant hotspot mutations and CGI methylations (22). Then, 4,269 mutations with p-values <=0.5 were selected from 17,313 hotspot mutations, and 5,277 CGI methylations with p-values <=0.05 were selected from 51,845 CGI methylations. Different p-value cutoffs were used to keep the feature counts comparable for the three data inputs as significance is an important filtering parameter. However, in the data preprocessing stage, it is more crucial to minimize the bias of distribution than emphasize the importance of features (23).

### Multiomics Multifactor Analysis and Description

To uncover the features associated with olaparib resistance, we classed the CCLE cell lines as either olaparib responders or nonresponders and then utilized multiomics factor analysis v2 (MOFA+, v2_1.2.2). The default training options for the model were used, including 15 factors and a fast convergence mode. To pinpoint the most significant features in each type of omics data, we selected the factors with the highest variance from each feature.

To identify features related to olaparib resistance, we conducted unsupervised matrix factorization. This process allowed us to find high variance in specific factors through variance decomposition and examine the components within those factors. We were able to find one factor with the highest variance for each of the omics features and selected biomarkers based solely on these factors.

We performed principal component gene-set enrichment (PGSE) to find features and components that could affect the response to olaparib and explain the factors that define each component (24–26). We classed the PGSE results as either upregulated or downregulated pathways and investigated the differences between the two types of pathways. We found the top 20 pathways for each analysis and identified shared pathways and genes.

To find features that distinguish responders from nonresponders using minimal information, we selected the features with the highest weights from MOFA. We tested weight cutoffs from 0.1 to 0.8 and used features with weights greater than 0.8 for further analysis (27).

### Biomarker Candidates and Model Construction

We created three prediction models using machine learning methods to test the performance of the candidate biomarkers. The models used were SVM, RF, and SNN (15,28–30). We generated the prediction models using R (v.4.1.1) and Python (v.3.9.7). SVM (e1071; v1.7-9) and RF (RandomForest; v.4.7-1) were used with R, while SNN (TensorFlow; v.2.9.0) was used with Python. Data were normalized using MinMax scaler equation (1). Since we used six biomarker candidates, the input formula was as follows equation (2).

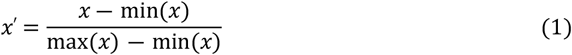

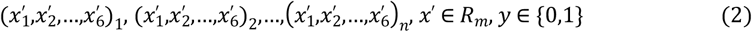

We used 70% of the 39 samples (27/39) as the training set. Y represents the category of the response and can be either 0 or 1. To resolve regression problems in the SVM model, we used support vector regression (SVR) equation (3).

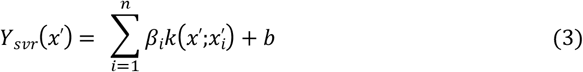

Support vectors are represented by β_i_ and x^’^_i_, and k is the kernel function. To prevent overfitting, we used the tune function and applied a sigmoid kernel for plane transformation equation (4).

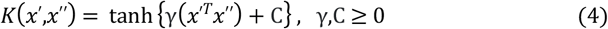

To optimize the hyperparameters, we used 10-fold cross validation for the hyperparameters γ and C, and the optimal parameters were found to be γ = γ = 10^-6^ and C = 10^5^. In the RF model, we used the same number of training samples as in the SVM model and evaluated the model by generating trees using bootstrap aggregating. We generated the bagging dataset using 500 trees from the n training samples (31) equation (5).

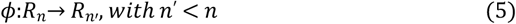

The total number of trees used was 500, and the error rate of the out-of-bag set was 34.21%. To estimate the marginal effect of each biomarker, the accuracy from the bagging dataset was used to measure the mean decrease in Gini impurity equation (6).

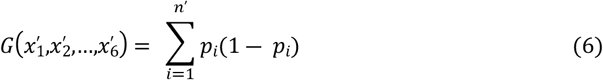

We generated the Siamese encoder, which includes all triplet data (*x*^′*A*^,*x*^′*P*^,*x*^′*N*^), by conducting random sampling. The Siamese encoder learned the difference between positive input x^’P^ and negative input x^’N^ with anchor input x^’A^. We created a structure that distinguishes responders from nonresponders by creating an encoding layer f through triplet training. We then mapped all the results to the latent space using the trained f. Finally, we added a step of binary classification by adding a classification model to the newly inferred data. We completed the model that distances the two classes with respect to a certain margin (α = 0.2) (16).

Anchor and positive inputs were extracted from the same category, while negative inputs were sampled from the other category. The training was performed so that the Euclidian distances between the anchor and positive inputs, |f(x^’A^) – f(x^’P^)|^2^, and the anchor and negative inputs, |f(x^’A^) – f(x^’N^)|^2^, are maximized and include the margin equation (7).

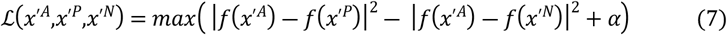

The Siamese encoder *f* was used with 11,137 parameters. The total number of epochs was set to 1000, with 200 steps per epoch and 20 validation steps. For X^’^ = (x^’^_1_,x^’^_2_,…,x^’^_6_), the mapping to the latent space E = e_1_,e_2_,…,e_m_ through *f* was used to add a fully connected layer for the binary classification of categorical data. The likelihood function L was determined by finding π that maximizes L(π|E), and the classification was performed equation (8). A fully connected layer was created and used with 279 parameters. The number of epochs used was 3,000.

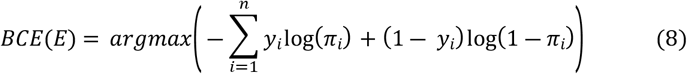

### Model Evaluation

We divided datasets into training datasets at 70% and test datasets at 30% to check the accuracy of the results. We calculated precision and recall for the three models and measured the F1 score and accuracy. We combined the individual models, generated SVM, RF and SNN, into a predictive model that predicts the response to olaparib using cell lines and evaluated its accuracy using the GDSC dataset (ANNOVA-fitted dose-response; v.25Feb20). We tested the results of the three models by distinguishing between olaparib responders and nonresponders based on cell-line viability results. We used data from 2,614 cell lines that measured cell viability in the presence of olaparib in the entire dataset and, considering data imbalance, determined the cutoff for olaparib-sensitive cell lines as cell lines with a ln(IC_50_) values less than or equal to 2 among 0.5, 1, 2, and 3 and determined nonsensitive cell lines as those with a ln(IC_50_) value greater than or equal to 5.5 (32).

## Results

### Public Data Processing

We investigated various omics features that can explain the refractory nature of olaparib responses by comparing data from cancer cell lines generated during the preclinical evaluation of the drug (206162Orig1s000). Using the CCLE datasets, which allow for quantitative analysis, we selected 19 responders and 33 nonresponders; we used undersampling to account for data imbalance. The criteria for selecting nonresponders were based on the classification of cancer types, resulting in the use of cell lines from ten breast cancers, six ovarian cancers, and three colorectal cancers (S1 Appendix Table S1). We included the OV-90 cell line, which was not part of the preclinical evaluation, as it was used in previous studies (33).

### Preprocessing and MOFA Analysis

We used RNAseq gene expression data, reduced representation bisulfite sequencing (RRBS) DNA methylation data, and merged mutation calls from the CCLE. The gene expression data were based on 51,611 transcripts, the mutation calls used 17,313 coding genes that were germline filtered and processed, and the RRBS data were based on 51,845 CpG islands in upstream genes. We filtered the data using highly variable features recommended by the MOFA developers, selecting only genes with a similar trend within the in-group but with statistically significant differences between the groups (https://biofam.github.io/MOFA2/faq.html). For the mutation datasets, we included at least 635 genes with two or more mutations. For the gene expression dataset, we analyzed differentially expressed genes and selected 3,295 transcripts using edgeR for statistical tests (21,34) with a p-value <0.1. For the selection of GCIs that showed a statistical difference between the responder and nonresponder groups, we performed the Wilcoxon rank-sum test and selected 2,361 CGIs with p-values <=0.05.

We utilized gene expression data to generate regulon information to infer the mode of action of olaparib responders and nonresponders (20). The algorithm we used, DoRothEA, infers the relationships between transcription factors and downstream genes. It uses four levels of evidence: 1) evidence obtained from a literature search, 2) direct interaction information obtained from ChIP-seq data, 3) information from other databases obtained through the *in silico* prediction of the mode of action, and 4) reverse-engineered results obtained using gene expression data. We used data from 289 proteins from the 1,396 transcription factors that could be calculated using results at level C or higher that included at least level 1) or level 2) evidence.

### Latent Factors of Olaparib Resistance

We used 3,296, 2,361, 636, and 289 features for gene expression, level of DNA methylation, deleterious mutation, and protein activity, respectively (Figure 2b). Each set of features was then matrix-factorized into nine factors (27). Among the four domains, deleterious mutations had the highest variance for the most dominant factor, while gene expression had the lowest variance. Although the sum of variance for protein activity was higher than that of gene expression, the pattern of variance was similar to that of gene expression (Figure 2c). The sum of variance for each domain was highest for mutations, followed by protein activity, gene expression, and DNA methylation (Figure 2d, S1 Appendix Table S2). To examine the differences between groups, we applied UMAP using latent factors (Figure 3) (35). The comparison of between-group differences using latent factors was more evident than when gene expression alone was used (Figure 3a). Most of the nonresponders were breast cancer cell lines (11/19), and the most common cell line in the responder group was colorectal cancer cell lines (5/19).

**Figure 1.**
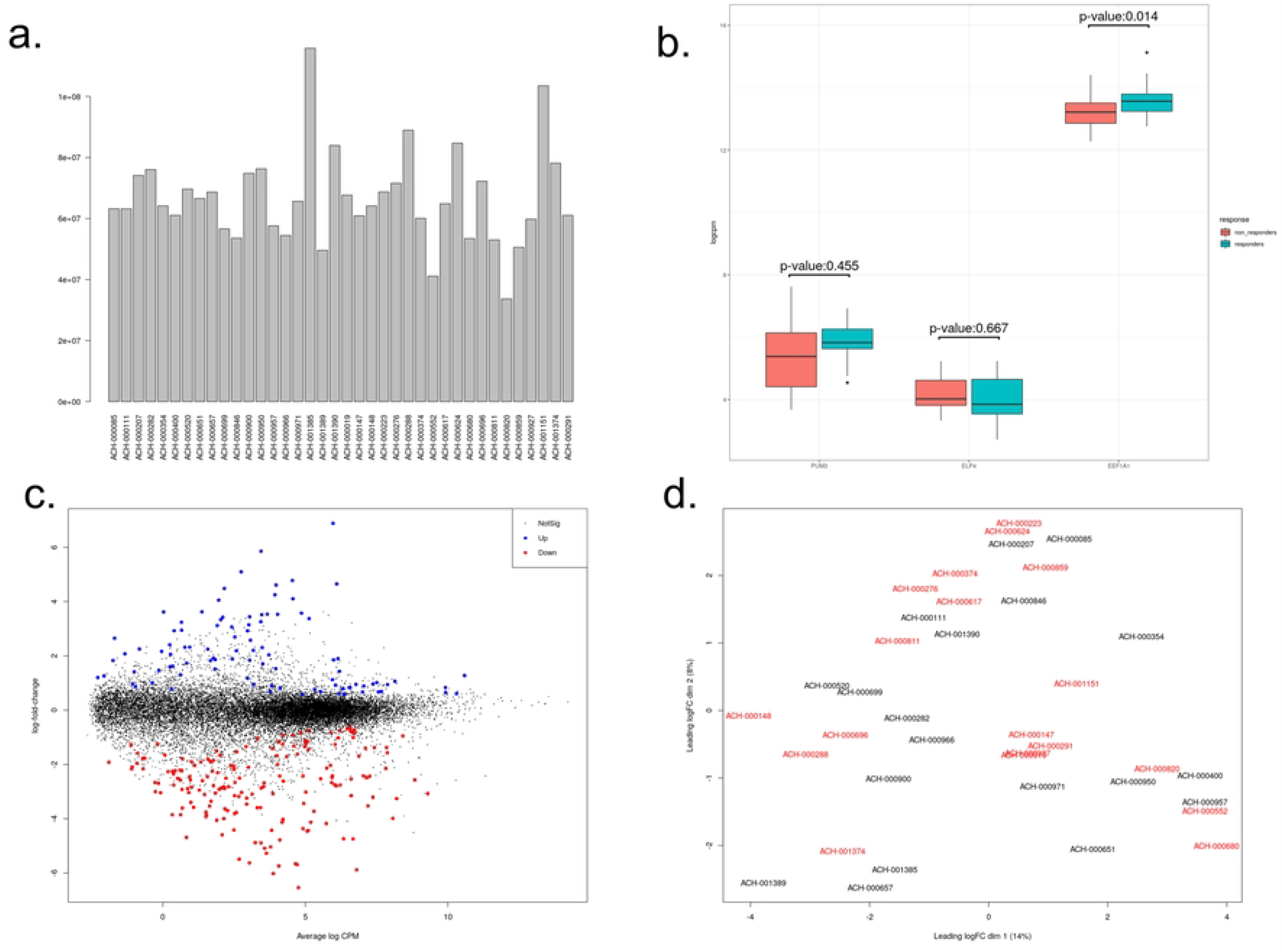
Gene Expression Quantification Using CCLE Database. a. Number of Read Counts for the Cell Lines Used. b. Evaluation of Results for Biomarker Identification. c. Genes with Significant Changes in Expression in Non-Responders. d. Mean-Difference Plot of Read Count Data.

**Figure 2.**
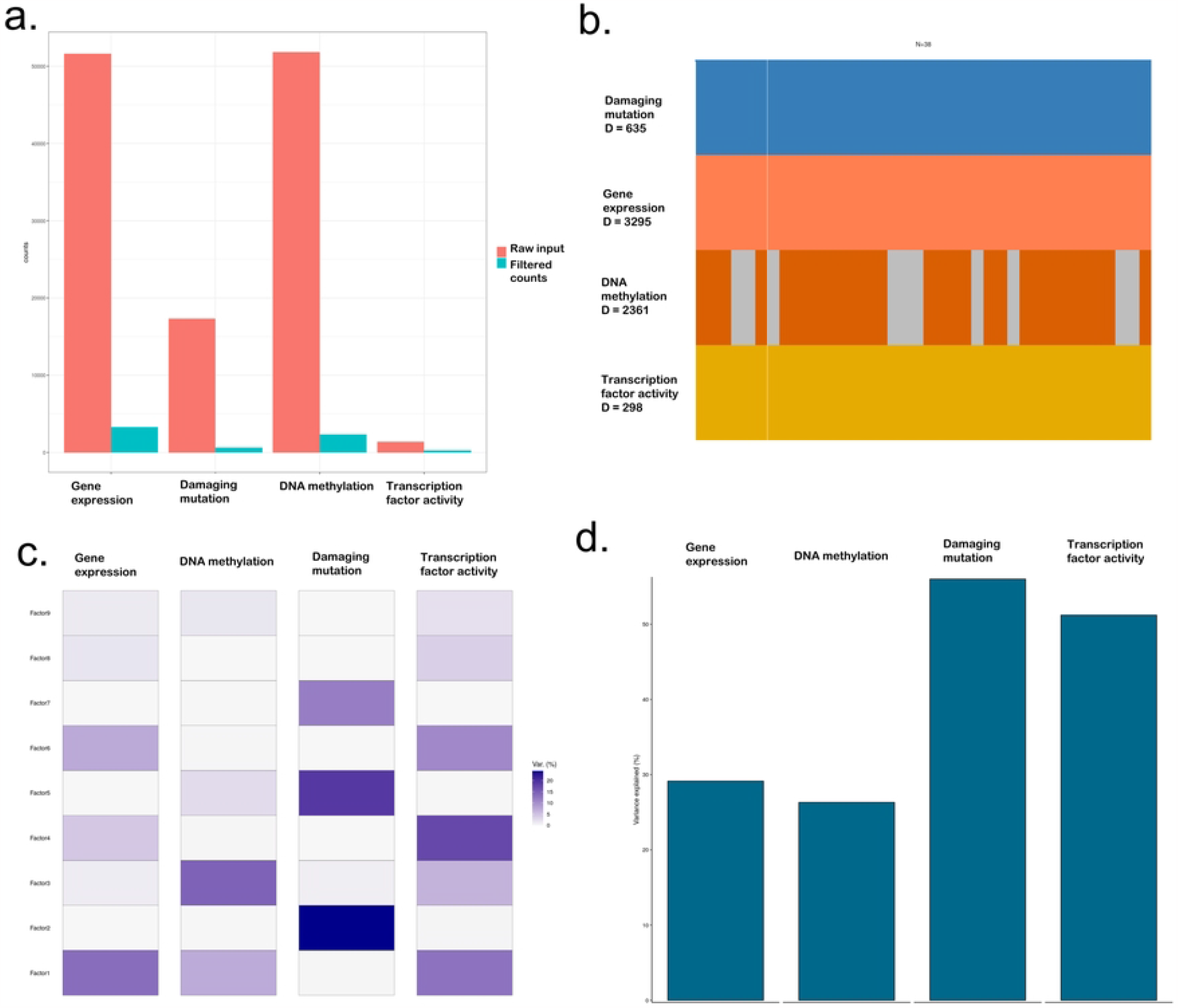
Overview of MOFA analysis results and factor variance. a. The number of data counts of filtering on all features. b. The number of cell lines and multi-omics features used as input. c. Calculated variance results for each factor of the four omics features. d. Sum of factor variances for each feature.

**Figure 3.**
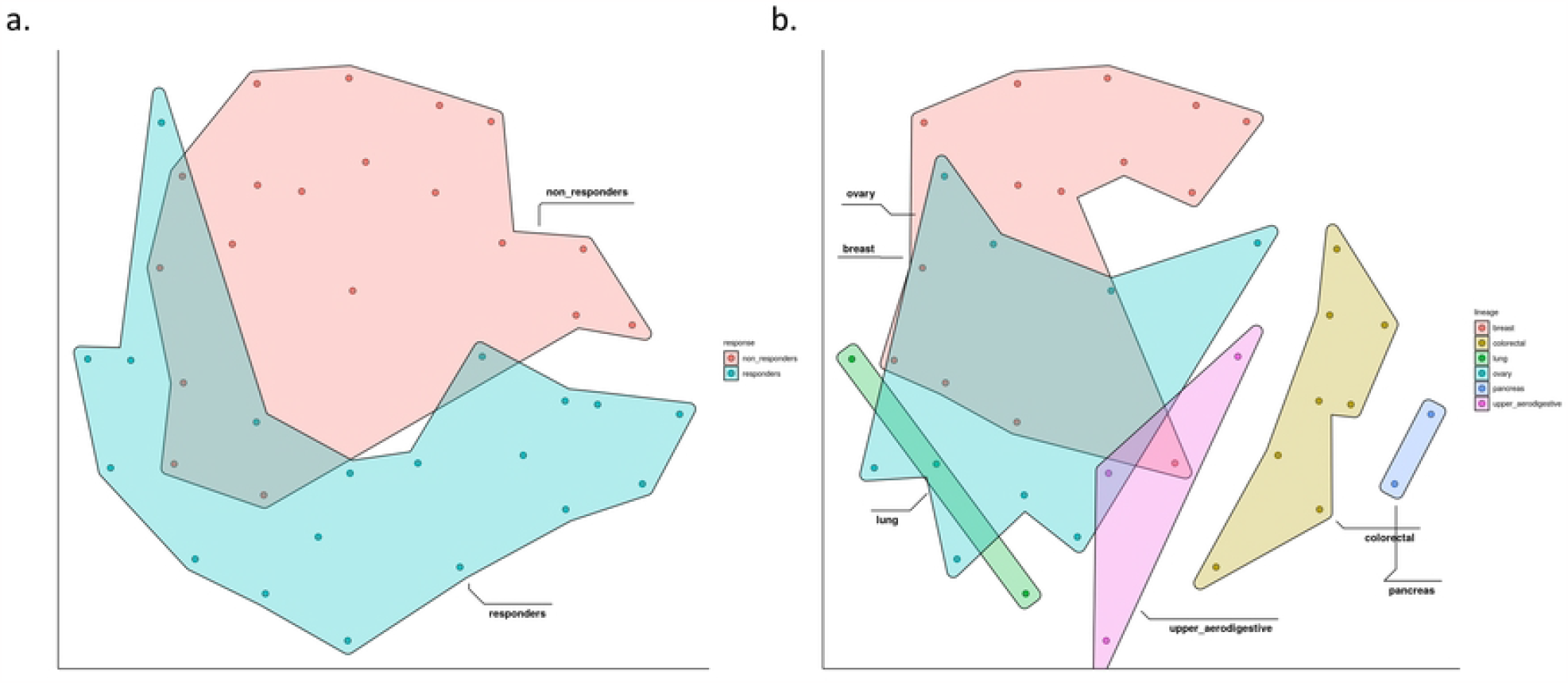
The UMAP plot is based on the latent factors obtained from MOFA analysis. It was clustered to show grouping based on two different factors: a. olaparib response and b. Cancer lineage.

### Reactome Analysis

We aimed to identify the molecular mechanisms that explain the differences between responder and nonresponder groups using gene expression data (25). The pathways that were more highly activated in the nonresponder group included extracellular matrix organization, collagen formation, and integrin–cell surface interaction (S1 Appendix Figure S3a). In the responder group, the pathways most highly activated were extracellular matrix organization and collagen formation as well as nonintegrin membrane ECM interactions (S1 Appendix Figure S3b). Both groups showed high activation of pathways related to the cell surface. Although some pathways were activated in both groups, the genes that mediated the pathway activation were different (S1 Appendix Figure S3c, d). The pathway components with the highest factor weights in nonresponders were LOXL2, SDC2, ADAMTS1, COL6A2, and FBN2 (S1 Appendix Figure S3c). LOXL2 is associated with the angiogenesis of primary breast cancer tumors, and inhibition of LOXL2 reduces angiogenesis activity (36). SDC2, a gene that encodes for syndecan, is associated with cancer progression through interaction with cell-cycle proteins and extracellular matrix proteins (37). It is also associated with multidrug resistance (38). ADAMTS1 is a member of the gene family that codes for secreted zinc-dependent metalloproteinases and interacts with vascular endothelial growth factor A (39). This gene is associated with breast cancer metastasis, and gene upregulation causes PPARδ activity to be suppressed, which helps prevent the invasion and migration of cancer cells (40). The pathway components with the highest factor weights in the responder group were P3H2, NTN4, SERPINE1, COL4A1, and ITGB3. SERPINE1 is directly related to drug resistance (41). Our reactome analysis indirectly shows that olaparib resistance is related to genes linked to extracellular matrix formation and function.

### Biomarkers of Olaparib Resistance

Using information obtained from reactome analysis, we next attempted to find biomarker candidates that could directly distinguish between olaparib responders and nonresponders (Figure 4) (23). We found 17, 6, 17, and 25 biomarker candidates in DNA mutation, gene expression, DNA methylation, and protein activity, respectively (S1 Appendix Table S2: weight value >= 0.8). For the 17 mutation marker candidates, which included IRS1, ZNF365, ARID4B, SLC12A9, CHRM3, ESRP1, CCDC15, ADNP, DDX27, GLYR1, NBEA, SBNO1, RPL22, ARSJ, ACACB, FSIP2, and ACVR2A, we conducted a gene-set enrichment analysis (42,43).

**Figure 4.**
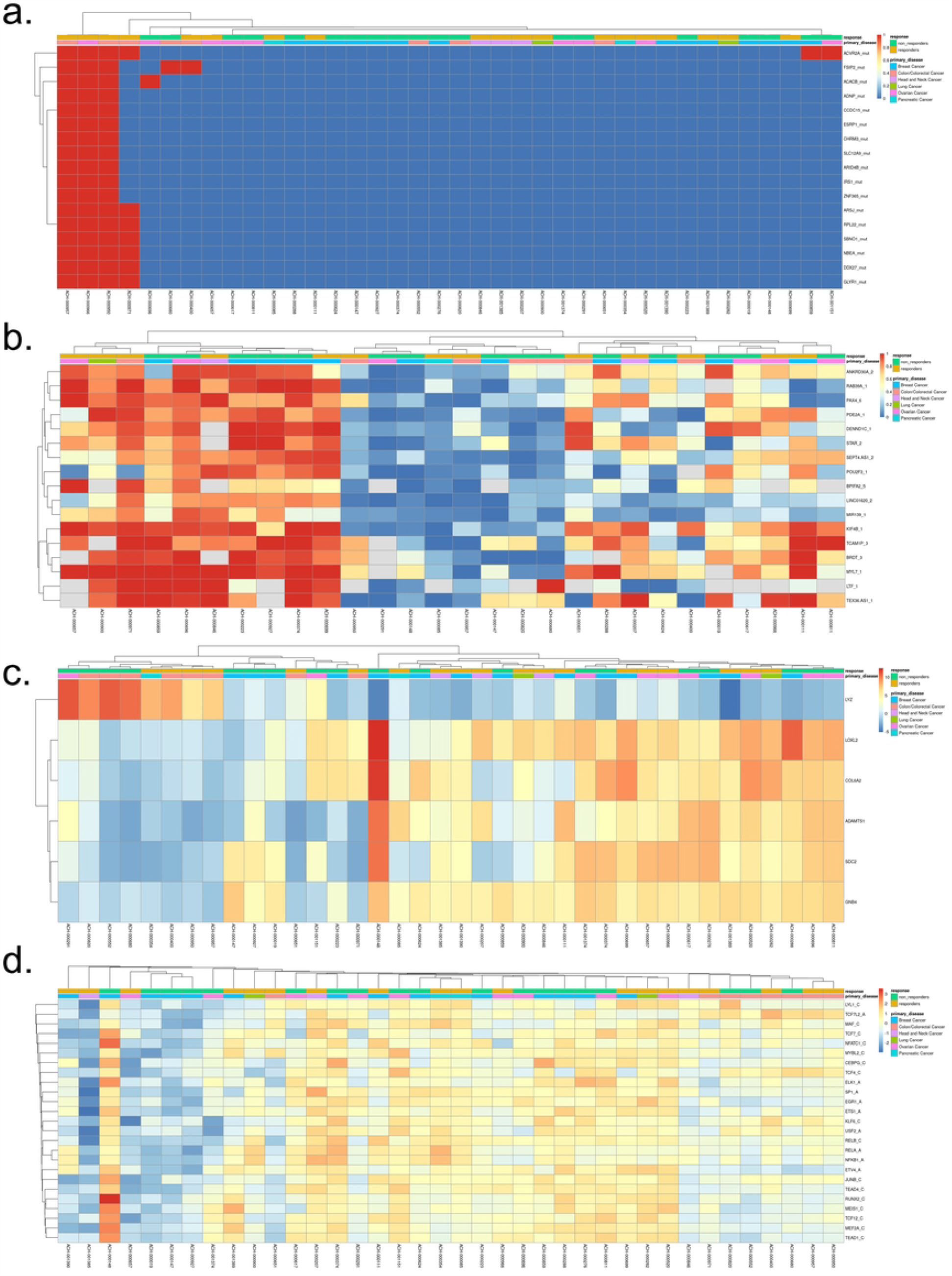
Quantification heatmaps of biomarker candidates. The first column on the top horizontal axis distinguishes between responders and non-responders, and the second column indicates the primary cancer type. a. Quantification heatmap using mutation markers, where a damaging mutation is marked as 1 and no mutation is marked as 0. b. Heatmap using CGI methylation level, where the left 10 cell lines are hyper-methylated compared to the right 18 cell lines. c. Gene expression quantification heatmap using the log(cpm) value from CCLE. d. Heatmap representing transcription factor activity using the DoRothEA score.

The gene-expression marker candidates LYZ, LOXL2, SDC2, ADAMTS1, COL6A2, and GNB4 showed the most significant functions in the extracellular matrix cellular component, The DNA methylation marker candidates were PDE2A, DENND1C, KIF4B, RAB39A, BRDT, TCAM1P, PAX4, POU2F3, BPIFA2, LINC01620, MYL7, STAR, LTF, TEX36.AS1, ANKRD30A, MIR139, and SEPT4.AS1. The protein activity marker candidates were ETS1, TEAD4, MYBL2, JUNB, KLF6, USF2, RUNX2, TCF4, SP1, RELA, RELB, CEBPG, MEF2A, NFKB1, NFATC1, TEAD1, TCF7L2, EGR1, TCF12, ETV4, TCF7, MEIS1, MAF, ELK1, and LYL1.

In the quantification of damaging mutations using mutation markers, most of the mutations were discovered in the responder group. Four cancer cell lines (ACH-000950, LOVO; ACH-000957, LS180; ACH-000966, IGROV1; and ACH-000971, HCT116) with many mutations had approximately ten times the number of mutations than the average number of damaging mutations in all CCLE cell lines (an average of 67.77 mutations, in 654, 714, 588, and 864 genes in the LOVO, LS180, IGROV1 and HCT116 cell lines, respectively). The average number of damaging mutations in all responders was 185.26, while that in nonresponders was 90.28. ACH-000966, IGROV1 is an ovarian cancer cell line, and the other three cell lines are large intestine cancer cell lines. On average, responders had more mutations, and all four cell lines were outliers (S1 Appendix Figure S1). IRS1, ZNF365, ARID4B, SLC12A9, CHRM3, ESRP1, CCDC15, and ADNP had the same weight values. The mutations in the four cancer cell lines were damaging mutations, except for those in IRS1 and ZNF365 (S1 Appendix Figure S2a).

When quantifying DNA methylation level using markers found in CpG islands, we observed a distinction between hyper-methylated genes and hypomethylated genes. The ten cancer cell lines with promoter hyper-methylation were comprised of five responders and five nonresponders, and the cell lines with hypomethylated regions were comprised of ten nonresponders and eight responders (Figure 4b). The genes PDE2A, DENND1C, and KIF4B had the highest weight values for differences in methylation levels in their promoter CpG islands, and these genes had similar patterns of methylation. KIF4B was hypermethylated in several cell lines (S1 Appendix Figure S2b).

All the marker genes except LYZ had similar expression patterns in all the cell lines, and the difference among the cancer cell lines was that the left 15 cell lines and the remaining 23 cell lines were divided into two groups in terms of expression patterns. The left group consisted of eight responders and nine nonresponders, while the other group consisted of 13 responders and 10 nonresponders (Figure 4c). The LYZ gene with the highest weight value encodes lysozyme and is involved in protein binding (to identical proteins) and lysozyme activity (S1 Appendix Figure S2c) (44).

The DoRothEA-based transcription-factor activity scores were transformed into z-scores to enable easy pattern recognition. Based on the z-scores, it was possible to divide the cancer cell lines into two groups based on transcription-factor activity: the 11 cancer cell lines shown on the left of Figure 4d and the 27 cancer cell lines shown on the right of Figure 4d. ACH-000148 (Hs 578T) is a breast cancer cell line with a relatively small number of damaging mutations (21 mutations), but it had higher protein activity than other cancer cell lines (Figure 4d). The gene ETS1 had the highest weight value and has been previously shown to be overexpressed in the breast cell line MCF-7, leading to the development of drug-resistant cells (45). In the DoRothEA data classification stage, ETS1 was the gene with the most evidence information and was rated as an ‘A-grade’ gene. ETS1 is a transcriptional activator or repressor and is related to stem-cell development, cell senescence and death, and tumorigenesis (46).

### Model Evaluation

We created a predictive model to explore whether the biomarker candidates found using MOFA could be applied to patient samples, and so we tested the ability of the biomarker candidates to predict drug resistance. We focused on gene expression data, which was the easiest genomic feature to access. We conducted tests to distinguish responders and nonresponders using a support vector machine (SVM) model and a random forest (RF) model to compare the performance of the predictive model with Siamese neural network (SNN). Firstly, we limited the number of biomarkers used based on the weight values obtained through matrix factorization. This allowed us to determine whether the performance could be maintained even if the number of genes decreases, and we also measured the efficacy of the biomarkers themselves. By utilizing SVM and RF, we confirmed how the prediction performance changes based on the number of features. We increased the weight values from 0.1 to 0.8 in increments of 0.1 and selected only the biomarker candidates with values higher than 0.8 (S1 Appendix table S3), and in the case of SVM, the precision values remained at 1.0 at all intervals. However, the recall value decreased, and the lowest performance was shown when the weight value was 0.7 (Figure 6a). In the case of the RF model, the precision value showed a trend of decreasing as the weight value increased, and the highest F1 score was obtained when only six features were used (Figure 6b).

We developed a predictive model to assess the efficacy of biomarker candidates by matching the existing drug-resistant label in the genomics of drug sensitivity in cancer (GDSC) database (47). To distinguish responders from nonresponders with limited data, we employed few-shot learning, which can be accomplished with SNN). SNN is a type of deep learning architecture where two identical networks are trained to compare the outputs of two different inputs (15,16). We utilized the six gene-expression biomarker candidates found from MOFA to train the model (Figure 5). To effectively differentiate responders from nonresponders, we minimized the unknown black-box part of the model and trained the base model using triplet loss. Then, we added a classification model to the latent vector that passed through the base model to determine if the latent vector of the six gene signatures belonged to the responder group or the nonresponder group.

**Figure 5.**
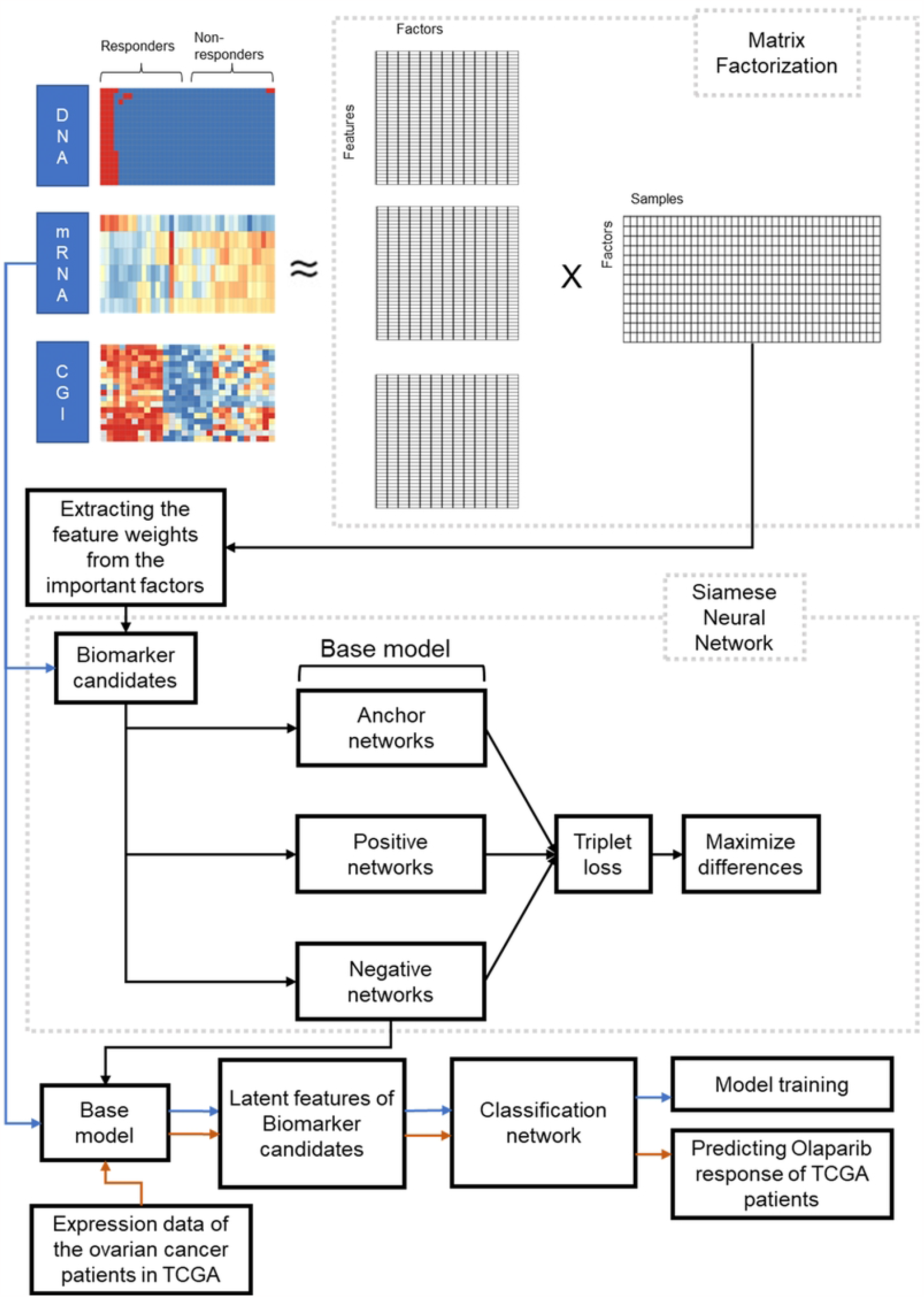
Figure 5 Workflow of a siamese neural network. This workflow consists of two main parts: selecting biomarker candidates by aggregating information on damaging mutations, gene expression, and DNA methylation, and performing classification using the selected biomarker candidates.

**Figure 6.**
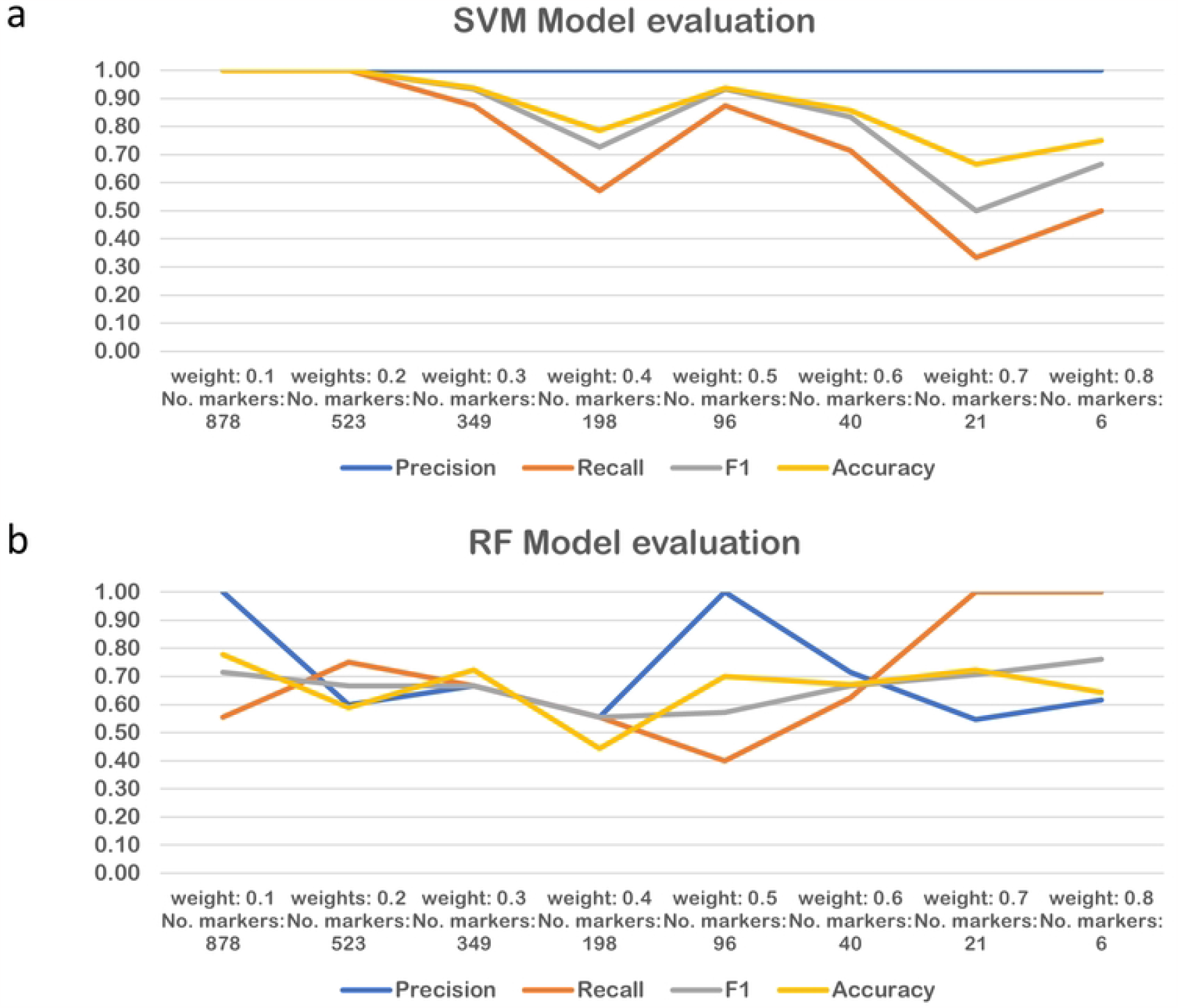
Performance comparison of SVM and Random Forest models with reduced marker count based on weights. a. Performance of SVM model. b. Performance of Random Forest model.

Evaluation of our predictive model showed that the SVM model had the highest precision value, the RF model had the best performance in recall, and the SNN model showed the highest performance in F1 score and accuracy (Table 1). This finding was consistent with the results of our experiments using preclinical evaluation results.

**Table 1.**
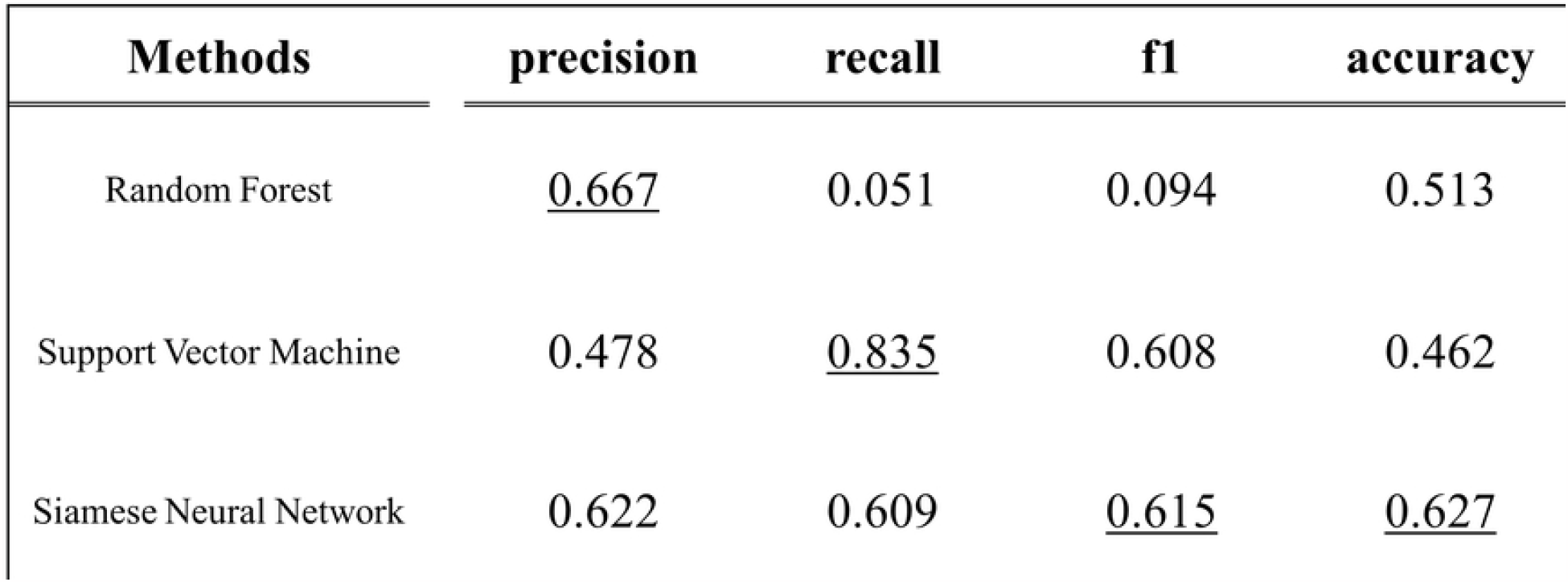
Assessment of the performance of the classification model on the GDSC dataset. The values underlined in the table correspond to the highest scores achieved for each evaluation criterion.

| Methods | precision | recall | f1 | accuracy |
| --- | --- | --- | --- | --- |
| Random Forest | <u>0.667</u> | 0.051 | 0.094 | 0.513 |
| Support Vector Machine | 0.478 | <u>0.835</u> | 0.608 | 0.462 |
| Siamese Neural Network | 0.622 | 0.609 | <u>0.615</u> | <u>0.627</u> |

## Discussion

Our study aimed to develop a predictive model for drug resistance using a combination of matrix factorization and machine learning techniques. We identified six potential biomarkers that were influenced by olaparib treatment using latent vectors obtained through matrix factorization. We applied these biomarkers to patient data and used survival information from the cancer genome atlas (TCGA) to categorize patients as responders, nonresponders, or others (S1 Appendix Figure S4). We treated outliers or noise arising from differences between gene expression signatures of cell lines and patients as a special class and classified to others (48). Our model showed significant differences in survival rates between responders (20%; 122/584) and nonresponders (10%; 62/584), indicating that the six biomarkers could be potential predictors of drug resistance.

However, our study had several limitations. We only used transcriptome data due to a lack of data and label information for other omics datasets. The Kaplan-Meier survival analysis showed that the two patient groups had significantly different survival rates; however, we did not have information on whether patients received olaparib. Further research is required to prove this (49). Additionally, the results of matrix factorization for the candidate biomarkers failed to clearly distinguish responders from nonresponders (Figure 3). Furthermore, the evaluation of our model using the GDSC dataset showed a decrease in performance compared to the existing dataset, likely due to the problem of dataset imbalance (Table 1).

Therefore, while our predictive model has potential applications in drug resistance research, it requires further validation through clinical trials before it can be applied to human experimentation. Our study also highlights the need for more comprehensive omics datasets and larger sample sizes to improve the accuracy of predictive models for drug resistance. Continued development of deep learning methods may also contribute to the creation of more accurate models with even less data.

## Conclusion

Our study demonstrates the potential utility of using a multiomics approach with machine learning techniques to identify biomarkers for drug resistance, as evidenced by the six gene-expression biomarker candidates identified for olaparib resistance; however, further validation through clinical trials and larger sample sizes are necessary to improve the accuracy and generalizability of the predictive model.

